# From spiral cleavage to bilateral symmetry: The developmental cell lineage of the annelid brain

**DOI:** 10.1101/268177

**Authors:** Pavel Vopalensky, Maria Antonietta Tosches, Kaia Achim, Mette Handberg-Thorsager, Detlev Arendt

**Affiliations:** Developmental Biology Unit, European Molecular Biology Laboratory, 69117 Heidelberg, Germany.; Max Planck Institute of Molecular Cell Biology and Genetics, Dresden, Germany.

**Keywords:** cell lineage, cleavage, spiralian, development, bilateral, symmetry

## Abstract

The spiral cleavage pattern is characteristic for Spiralia (Lophotrochozoa), a large assembly of marine invertebrates. In most cases, spiral cleavage produces freely swimming, trochophora-type larvae with a simple nervous system that controls ciliary locomotion. These larvae acquire bilateral symmetry, as manifested for example in the larval brain. The transition from the rotational symmetry of spiral cleavage into the bilateral adult body has not yet been understood. Here, we present the developmental cell lineage of the brain of the annelid *Platynereis dumerilii* from the zygote until the mid-trochophore stage (~30 hpf), in combination with a gene expression atlas for several embryonic and larval stages. Comparison of multiple embryos reveals a highly stereotypical development and an invariant cell lineage of the differentiated cell types. In addition, we observe a fundamental subdivision of the larval brain into a highly proliferative dorsolateral region and an early differentiating ventromedial region that gives rise to the apical nervous system. The transition from rotational to bilateral symmetry progresses gradually from the lateral to the central regions. Strikingly, the spiral-to-bilateral transition does not involve extensive cell migration. Rather, corresponding cells in different spiral quadrants acquire highly divergent identities in line with their bilateral position.

## INTRODUCTION

During early development, embryonic cleavage produces blastomeres via a rapid series of cell divisions without significant growth, relying on maternally deposited messengers and proteins. During these processes, the initially broad developmental potential of blastomeres becomes gradually restricted towards distinct cell fates. This can occur via two basic modes, called regulative (conditional) or mosaic (determinate) development. In regulative development, exhibited by vertebrates, cnidarians and sea urchins (Gilbert, 2000), almost all blastomeres share a broad developmental potential and cell fate determination largely depends on local signaling events. In mosaic development, most blastomeres inherit distinct maternal determinants and signaling is assumed to play a minor role. This requires sophisticated *in ovo* localisation, a stereotypic arrangement of cleaving blastomeres and an invariant cell lineage. Prime examples for mosaic development are the nematodes (Sulston et al., 1983) and ascidians (Nakamura et al., 2012; Nishida, 1987). Recent results however hint at a considerable degree of cell-cell signaling in their invariant lineage (Lawrence and Levine, 2006; Lemaire, 2009), which underscores that regulative and mosaic development only differ in the relative contributions of autonomous versus conditional cell fate determination.

Mosaic development with invariant early cell lineage and autonomous cell fate decisions is also characteristic for the Spiralia, or ‘Lophotrohozoa’, a large assembly of marine invertebrate phyla (Henry, 2014). Their eponymous ‘spiral cleavage’ produces a highly stereotypic, spiral-like arrangement of blastomeres (Fig. 1A) (reviewed in (Nielsen, 2004, 2005): The first two cleavages, perpendicular to each other, subdivide the embryo along the animal-vegetal axis into four blastomeres, representing the four future embryonic ‘quadrants’ A, B, C and D (Henry, 2014). The subsequent cleavages are asymmetrical, generating quartets of smaller micromeres towards the animal pole and quartets of bigger macromeres towards the vegetal pole. In addition, due to an oblique angle of these divisions, the originating micromere quartets are alternately turned clock- or counterclockwise against the macromere quartet, so that the micromeres come to lie in the furrows between the macromeres (Fig. 1A). This way the characteristic spiral-shaped organization of the embryo is generated. The initial cleavage pattern is identical for each quadrant, so that the whole early embryo shows rotational symmetry around the animal-vegetal axis. Corresponding cells with similar lineage in the four quadrants are referred to as *quadriradial analogues.*

**Figure 1:**
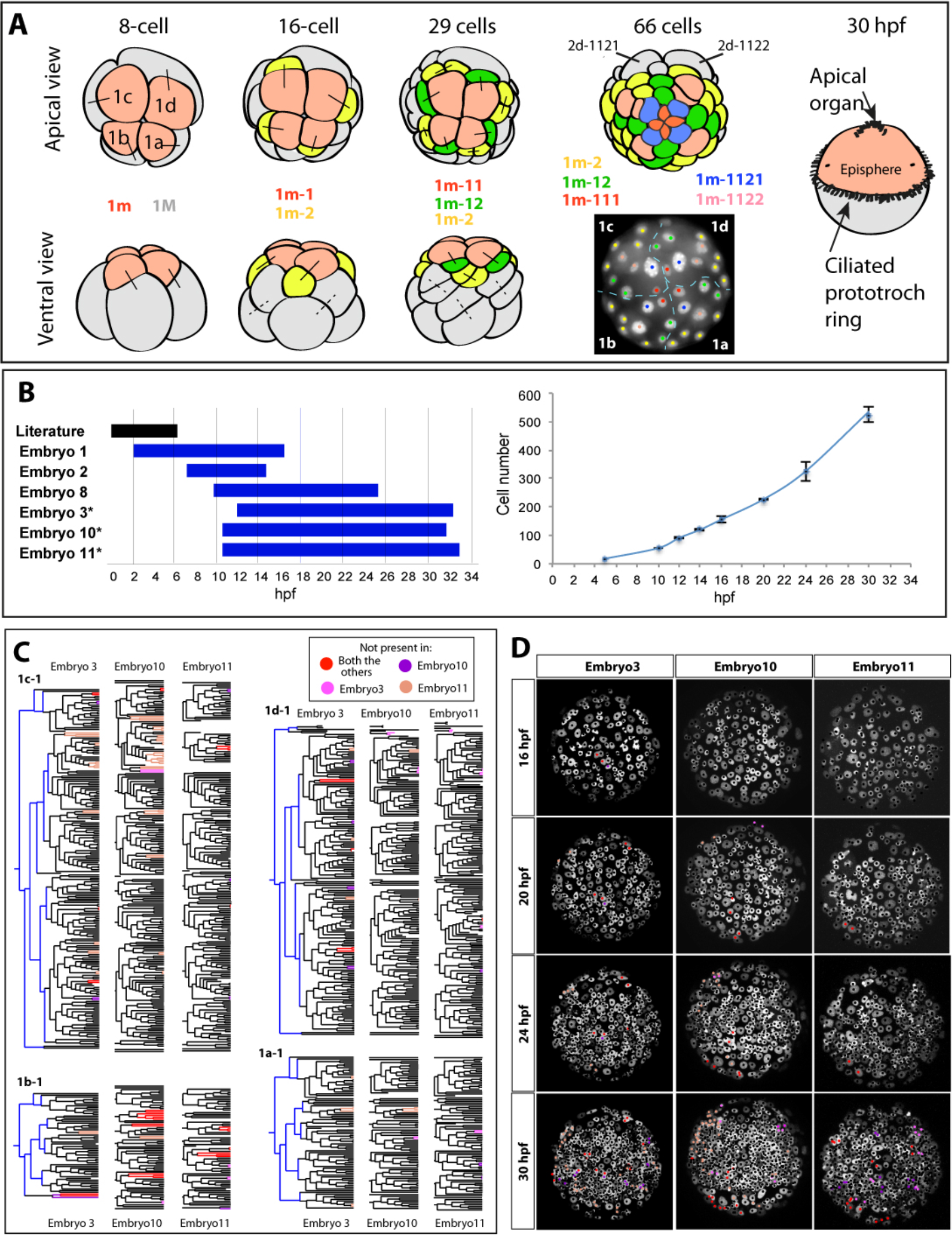
Life-imaging, tracking and comparison the cell lineages of the developing micromeres. (A) The early development of a prototroch larva by spiral cleavage. The apical quartet of micromeres 1m (light red) present at 8-cell stage gives rise to the episphere. The precurosrs (1m-2) of the prototroch cells are labeled in yellow. Apical views modified from (Dorresteijn et al., 1993). The ventral views are extensively schematized for simplicity. For 66-cell stage a schematic apical view (top) together with a snapshot (bottom) of the time-lapse recording of the developing episphere is shown. The color-coding of the nuclear tracks in the snapshot corresponds to the coloring in the schematic apical view. The dashed blue line represents the border between embryonic quadrants. (B) The overview of time-lapse movies used for the analysis. Nuclei counts of at least three fixed specimens for each stage were used for temporal calibration of the movies. Asterisks mark the movies used to create the consensus lineage tree. (C) The comparison of the cell lineage trees of three larvae up to 30 hpf. The early lineage covered in previous work but not covered by the movies is shown in blue. The corresponding cells/divisions conserved in all three larvae are colored in black. The divisions and cells that are not present in all three larvae are color-coded according to the legend. (D) The snapshots of the z-projection of the life imaging movies showing the differences between the three larvae at different time points. Differences are color-coded as in (C).

In most spiralian phyla, spiral cleavage produces spherical planktonic larvae with characteristic apical ciliary tuft and bands of motile cilia that drive larval swimming. In annelids and molluscs, these are called trochophora larvae (Fig. 1A). The larvae form a simple nervous system that integrates sensory information from photo-, mechano- and chemosensory receptor cells for the control of ciliary locomotion (Jekely et al., 2008; Marlow et al., 2014; Tosches et al., 2014). Its most prominent features are an apical nervous system with an apical organ underlying an apical tuft. The apical nervous system is connected via radial nerves to a ring nerve (Nielsen, 2004, 2005). The ring nerve innervates a pronounced circular ciliary band, the prototroch, which subdivides the larva into upper episphere and a lower hyposphere. During settlement metamorphosis, the larva transforms into an adult body with overt bilateral symmetry (or more or less complex derivatives thereof, see for instance the development of Crepidula (Hejnol et al., 2007; Lyons et al., 2017) and Ilyanassa (Goulding, 2009). The adults of most phyla develop a worm-shaped body form as manifest for example in basal mollusks and, most prominently, in flatworms and annelids. In these worms, the former episphere of the larva develops into the head including the paired cerebral ganglia. The hyposphere gives rise to the trunk including the ventral nerve cords (Nielsen, 2004, 2005).

Hence, the most peculiar feature of spiralian development is the transition from rotational (spiral) point symmetry to bilateral symmetry, which has puzzled embryologists for more than a century (e.g., (Wilson, 1892)). How is this spiral-to-bilateral transition accomplished? In the hyposphere, bilateral symmetry is established through the unique behavior of two cells, 2d and 4d (Wilson, 1892), which derive from the D-quadrant. These cells divide once into the left and right bilateral founder cells of the entire trunk, which proliferate in a stem cell-like fashion and give rise to trunk ectoderm and mesoderm, respectively (Fischer and Arendt, 2013; Gline et al., 2011; Lyons et al., 2012). The situation is more complicated in the episphere, where the bilateral symmetry has to emerge from a pre-existing array of spirally arranged micromeres. Here, the spiral-to-bilateral transition may involve a ‘rearrangement’ of micromere position via complex cellular movements, or start from selected bilateral founders analogous to the trunk founders. The latter solution was favored by E. D. Wilson (Wilson, 1892), who gave an early and detailed account of spiral cleavage in the annelid *Nereis*. He observed a sudden transition from spiral to bilateral cleavage pattern after the appearance of the prototroch that he attributed to such (yet to be identified) founders.

To decide between these options, we reconstructed the full developmental cell lineage for the marine annelid *Platynereis dumerilii* from the fertilized egg to the swimming trochophore stage (~30 hpf), by confocal microscopy. We linked early lineages to gene expression using a cellular resolution gene expression atlas for several embryonic stages (compare (Vergara et al., 2017)). This resource is explored here for the episphere spiral-to-bilateral transition. *Platynereis* is especially suited to study this transition, because the larval phase is rather short and the bilateral head and trunk structures are developed in the swimming trochophore, so that the full transition can be observed microscopically. Previous studies in *Platynereis* and other spiralians had established the bilateral fate of early micromeres by injection of tracer dyes, yet did not resolve their lineage in cellular resolution (Ackermann et al., 2005; Hejnol et al., 2007; Meyer et al., 2010). The time-lapse recordings, software tools and lineage analysis presented here generate an unprecedented resource for spiralian biology available so far only for nematode and tunicate model systems.

Our lineage analysis allows tracking the spiral-to-bilateral transition in cellular detail. As postulated by Wilson, we identify bilateral founder cells; yet, we observe a whole array of bilateral founders distributed over the episphere at around 12hrs of development. Some of them, located in the lateral episphere represent quadriradial analogues, that is, they stem from similar (i.e., corresponding) lineage in their respective quadrants. Others, located more medially, stem from dissimilar lineage in their respective quadrants. Mapping the expression of the conserved bilaterian head patterning genes *otx* and *six3* onto the developmental lineage, we find that lateral *otx* expression marks the bilateral founders with similar lineage, whereas medial *six3* marks those of dissimilar lineage. Moreover, we find that while the *otx+* lateral founders show strong proliferation during larval stages and remain mostly undifferentiated at 30hpf, the *six3+* medial founders differentiate earlier and give rise, among others, to bilateral pairs of cholinergic neurons in the larval brain. Finally, we find that the apical organ proper does not derive from bilateral founders at all. Instead, it develops from the most medial cells that do not show bilateral symmetry.

We discuss our findings as the result of a secondary overlap of two initially separate phases of development, namely an early mosaic and determinate and a later regulative phase, representing larval and adult development, respectively.

## RESULTS

### Live-imaging and tracking of the cell lineage in the *Platynereis* episphere

The brain is almost entirely formed by the offspring of the apical micromeres 1a-1d, collectively referred to as “1m” (Ackermann et al., 2005; Dorresteijn, 1990), easily accessible to life imaging by standard scanning confocal microscopy. To track cell divisions in the apical brain, we injected embryos at different stages post fertilization (1, 2 or 4 cell stage) with H_2_B-dsRed and Lyn-EGFP mRNAs (Haas and Gilmour, 2006), which label chromatin and cell membranes, respectively. Then we recorded time-lapse movies of these apically mounted embryos (Fig. 1A-B, Supplementary package 1). To track and reconstruct the lineage, we developed a package of simple macros for ImagejJ/FIJI allowing manual tracking and visualization of lineage-related information from confocal microscopy stacks. The package and documentation are available in the supplementary material (Supplementary package 1). We tracked all cell divisions in the episphere of multiple embryos spanning the developmental time from 16-cell stage (~2 hpf) until ~32hpf at which stage more than 500 cells are present in the episphere (Fig. 1B), with at least three embryos coverage per developmental stage. This comprehensive dataset allowed us to perform detailed cell lineage analyses of developmental stereotypicity, clonal behavior and the transition from spiral to bilateral symmetry.

### The cell division patterns of the *Platynereis* episphere is stereotypical at swimming larval stages

To challenge the stereotypicity of cell division patterns, we injected nuclear tracers into 2− and 4− cell stage embryos and the compared resulting clonal domains with the results of the live imaging. These validation experiments indicated that the clonal domains originating from tracer dye injections were in a good agreement with the shape and position of the clonal domains inferred from the tracked time-lapse movies (Supplementary Fig. 1A-D’), pointing to a high level of stereotypicity. In addition, the shape and overall arrangement of the clonal domains originating from ~13hpf are highly similar between embryos (Supplementary Fig. 1E). To address the stereotypicity of episphere development beyond this time point, we used an algorithm that identifies automatically corresponding cells in different movies on the basis of lineage information, relative cell positions at division and cell cycle length (Supplementary methods and Supplementary Fig. 1F-J). We compared the time-lapse movies of more than four independent embryos up to 24hpf and three embryos until 30hpf (Fig. 1C-D). This comparison revealed a largely stereotypical development, both at the level of the lineage tree topology as well as cell positions, with only a small number of differences distributed over the embryo (Fig. 1C-D). The embryos show no differences until 16hpf. The only exception is the ventral apical rosette cell 1b-111, which shows most variability in timing and division pattern observed across larvae (see below). Taken together, our comparative analysis shows that *Platynereis* brain development is highly stereotypical at the level of overall cell arrangement and lineage tree topology.

### The cell lineage of the first differentiated cells is invariant

Although the stereotypic tree topology and cell positions suggest an invariant cell lineage, we wanted to prove this assumption by investigating whether the same cell types are produced by the same cell lineage in different embryos. To date, the only differentiated cells for which the cell lineage has been fully described in *Platynereis* are the primary prototroch cells (Dorresteijn, 1990). In our time-lapse recordings, several differentiated cell types could be directly identified based on their morphology and other microscopical features: the prototroch cells, larval eye pigment cells, the five ventral gland cells (Fig. 2A) and several cell types in the apical organ (Fig. 2B) described in (Marlow et al., 2014). In addition, we fully mapped the expression of the cholinergic marker *Choline Acetyl Transferase (ChAT)* on the lineage, by performing whole mount mRNA in situ hybridization (WMISH) on several embryos fixed just after the last timeframe of the recording (Fig. 2C). At 30hpf stage, the *ChAT* expression pattern comprises typically nine differentiated cells, mostly involved in controlling cilio-motor behavior (Jekely et al., 2008; Tosches et al., 2014). Another hallmark of differentiating neurons is the formation of axons. Zygotic injection of a nuclear marker mRNA followed by the injection of LA-EGFP mRNA (which labels actin filaments) into one blastomere at 2-cell or 4cell stage allows following the cell lineage of cells forming axonal projections (Figure 2D-E’). With this approach, we identified two apical cells projecting outside the AB-domain (Figure 2D-D”) and cells with axons traversing the dorsoventral midline (Figure 2E-E’).

**Figure 2:**
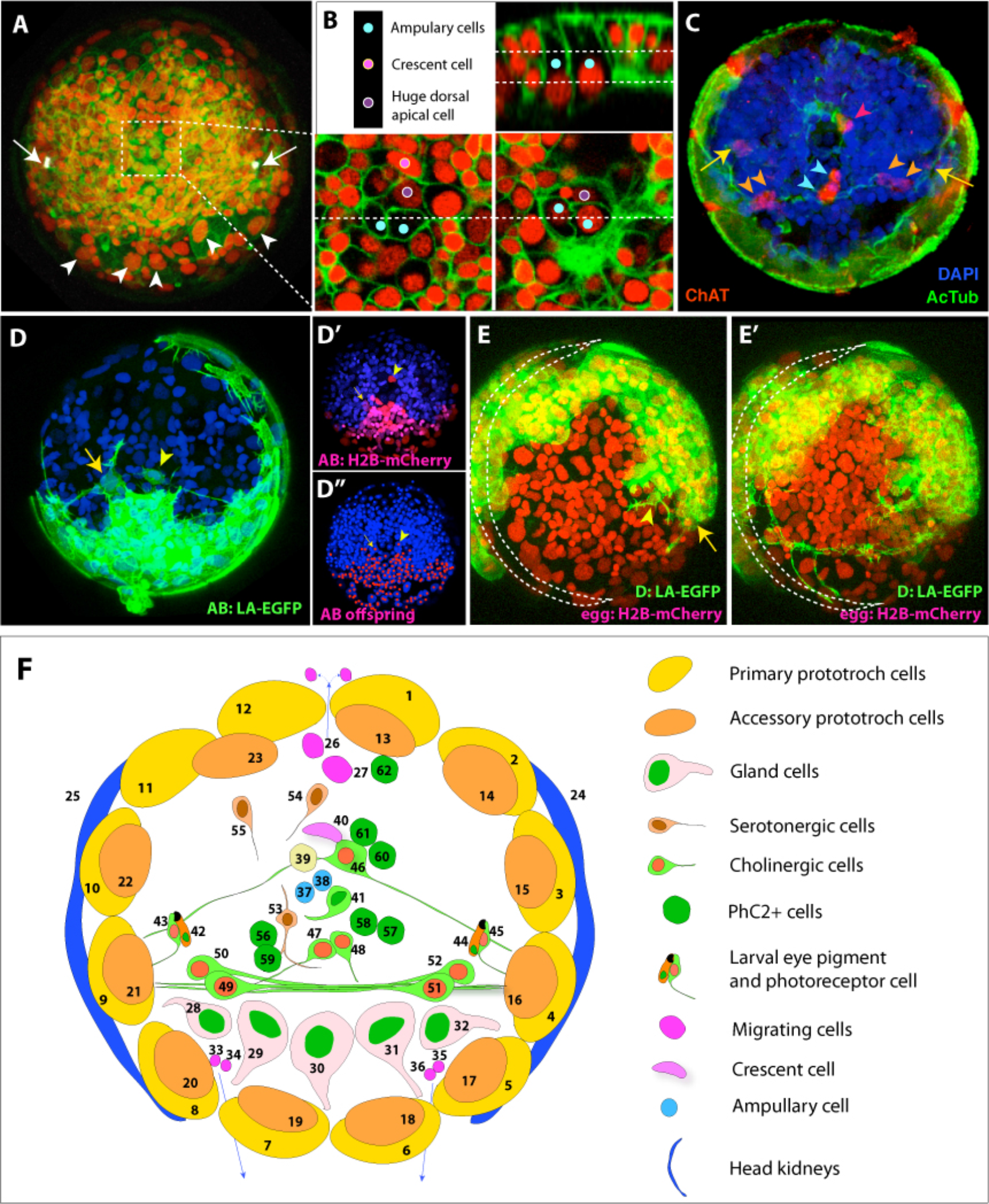
Differentiated cell types with known lineage at 30 hpf. (A-B) Several cell types identified directly in the last frame of the time-lapse recordings by their morphological features. The larval eye pigment cells identified by their autofluorescence in the red spectrum (A, white arrows). The large gland cells with the typical flask-shape (A, white arrowheads). (B) Several cell types of the apical organ could be identified by morphology and position (described in detail in Marlow et al, 2014). (C) The WMISH of ChAT performed on the live-imaged larva fixed immediately after imaging allows establishing the cell lineages of cholinergic neurons. Yellow arrows indicate the position of larval eye photorecpetors, orange arrowheads the ventrolateral ChaT+ cells, red arrowhead the apical ChaT+ cells with first lateral axons, and blue arrowheads the ventromedial cholinergic cells. (D) Two apicalneurons with axons revealed by injection of the AB blastomere with LA-EGFP mRNA. The neuron with the ventrolateral projections stands out of the rest of the AB labeled domain (D’ and D”). (E-E’) The snapshots of a time-lapse recording of larvae injected with H2B-mCherry mRNA at one cell stage and LA-EGFP to the D blastomere shows the axon of a flask-shaped cell in the apical organ (yellow arrowhead) and the growing axon of the ventral cholinergic cells (yellow arrow). The prototroch ring is indicated by a dashed crescent. (F) A summary diagram of the differentiated cell types in the episphere at ~30hpf. The numbering corresponds to the first column of Table I.

In summary, we addressed the cell lineages of 62 differentiated cells in a 30hpf episphere, summarized in (Figure 2F and Table I). For the vast majority of investigated cell types (exiting the cell cycle before ~15hpf), the cell lineage is strictly conserved among multiple embryos (column “Support” in Table I). Interestingly, the cell lineage varies substantially in later-born cells (ChAT-positive cell Nr 52 exiting the cell cycle at ~28 hpf, cell Nr 50, ~20hpf). Based on the analyzed embryos and available literature we created a consensus lineage tree with the identified cell types (Supplementary Fig. 2). In summary, our analyses show that the *Platynereis* larval brain develops via stereotypical cell divisions and that the lineage of early differentiating neuronal cell types is conserved.

**Table I:**
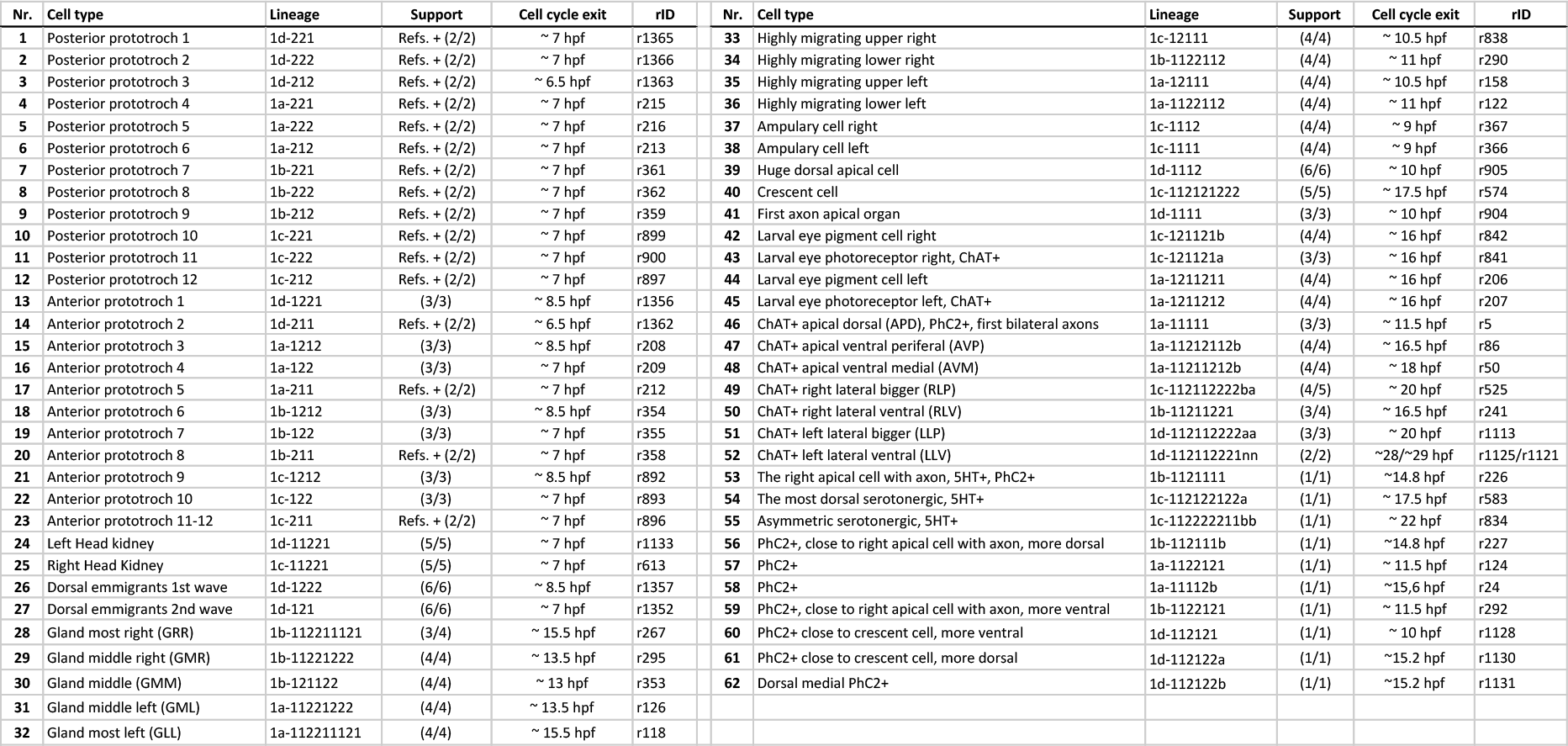
The cell lineage of the differentiated cell types at 30 hpf. The numbers in the first column correspond to the numbering in Fig. 2F. The column “Lineage” contains the consensus lineage based on the literature and multiple time-lapse recordings. The “Support” column indicates the number of time-lapse recordings in which the same lineage/the number of total time-lapse recordings analyzed for a given cell types. The “rID” column contains the reference ID of the given cell type which corresponds to the rID in the tracking files (Supplementary package 1) and the consenus tree (Supplementary Fig. 2).

### A gene expression atlas for embryonic and early larval stages

To link cellular lineage to differential gene expression, we conducted whole mount *in situ* hybridization expression analysis for a total of 23 genes for 6 stages (12, 14, 16, 20, 24, 30) (Supplementary Fig. 3 and 5). Since most of the identifiable early differentiating larval cells represent neural cell types, we focused on neural regionalization transcription factors such as the homeodomain factors *otx, six3/6, nk2.1,* neural specification factors such as the Group A basic Helix-Loop-Helix (bHLH) factors *ato*, *neuroD*, *ngn*, *ascl*, *ash2*; the Group B bHLH factor *olig*, the Group E bHLH factor *deadpan*, the Group F bHLH factor *coe* (*Supplementary Fig. 3A*), the homeodomain factor *prox*, *and* the C2H2 zinc finger *klf*, (*12-24 hpf, Supplementary Fig. 3B*); and between 12 and 30 hpf general neural differentiation markers such as *elav* and *synaptotagmin (syt)* and more specific markers such as prohormone convertase 2 *(phc2)* for neuropeptidergic cells (Tessmar-Raible et al., 2007), vesicular acetylcholine transporter *(vacht),* choline acetyltransferase *(chat)* for cholinergic neurons, vesicular glutamate transporter *(vglut)* for glutamatergic neurons, tryptophan hydroxylase *(tph)* for serotonergic neurons, and larval eye markers such as the neuropeptide *FVRI,* rhabdomeric opsin *r-opsin3* (Jekely et al., 2008), and pigment cell marker tryptophane-2,3-dioxygenase *(Trp2,3) (Supplementary Fig. 3C)*.

Using our atlas, we found that the transcription factors *coe*, *ngn*, *neuroD*, and *prox* are co-expressed with the neuronal differentiation markers *elav and Syt*, the cholinergic marker *ChAT*, and the neuropeptidergic marker *Phc2* in the apical organ cells (Nr. 46 and Nr. 53, later serotonergic, in Table I). At later stages, even when expressing cells could no longer be identified individually, our analysis revealed expression correlations and transcriptional dynamics in neural lineages. For example, the expression of the neuronal specification factors *prox*, *ngn* and *NeuroD* appears to faithfully anticipate expression of the pan-neuronal marker *elav* (compare Supplementary Fig. 3A-C). Similarly, we observed that expression of the bHLH factor *coe* precedes the expression of cholinergic markers *VAChT* and *ChAT* several hours later (Compare Supplementary Fig. 3A and 3C), in line with the evolutionary conserved role of COE factors in specification of cholinergic neurons (Kratsios et al., 2012). Interestingly, expression of the two neuronal differentiation markers *phc2* and *syt* remains restricted to the apical organ region between 24-34 hpf, partially overlapping with the cholinergic markers ChAT and VAChT. This suggests that cholinergic and neurosecretory cells form the core of the larval apical nervous system, in line with single-cell RNA sequencing results (Achim et al., 2018). The restricted and stable expression of *Phc2* and the cholinergic markers contrasts to a rather dynamic expression of *NeuroD*, *ngn*, *and elav* that demarcate neuronal specification more broadly in the developing cerebral ganglia.

### Lineages retaining rotational symmetry: prototroch and apical organ

We next used the tracked lineage of *Platynereis* embryos and larvae to analyse the symmetry properties of individual cellular lineages. In conjunction with the gene expression atlas, this allowed us to correlate symmetry properties with gene expression and cellular differentiation.

We first focused on lineages that retained the initial rotational symmetry. In *Platynereis*, these lineages give rise to early differentiating cells of the prototroch and apical organ. The primary prototroch develops from the two vegetal-most quartets of the first micromeres, that is, 1m-22 and 1m-21, in a strictly radial arrangement (Fig. 3A-B). The blastomeres 1m-12, located slightly more apically, divide twice in a spiral mode (with an exception of 1d-12, see below) (Fig. 3B). They produce the non-dividing accessory prototroch cells 1m-122 and 1m-1212.

**Figure 3:**
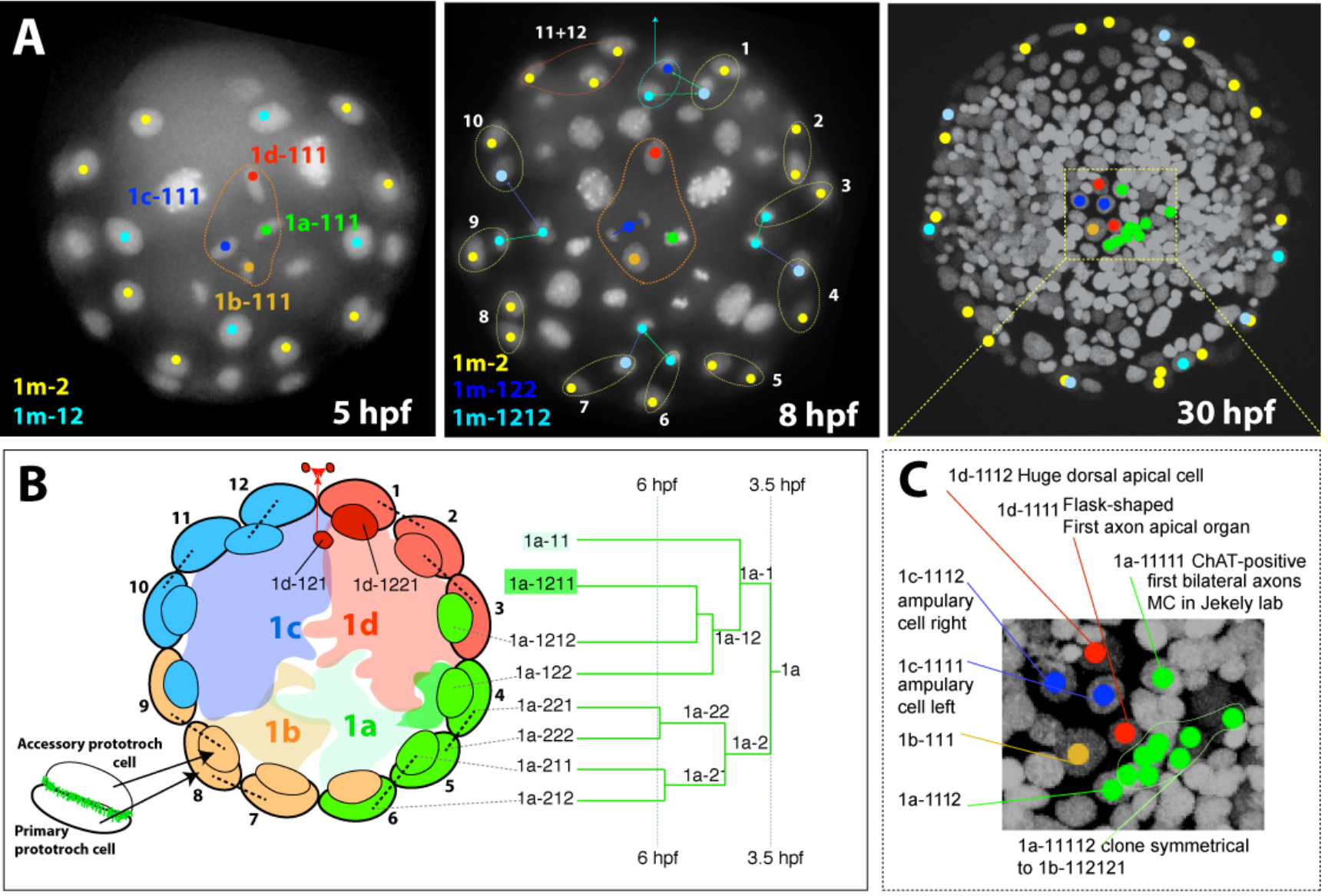
The developmental origin of the accessory prototroch cells and the cells in the apical organ. (A) The overview of the development of the apical rossette (1 m-111, orange dashed line) and the primary (1m-2, yellow) and accessory (1m-12, light blue) prototroch cells at 5, 8 and 30 hpf. (B) Schematic diagram showing the lineage origin of the prototroch cells. Only the 1a quadrant lineage tree is shown for simplicity. The cells in the scheme are color coded by their quadrant of origin and the black dashed lines indicate sister cells. Each quadrant contributes three primary prototroch cells (1m-222, 1m-1212 and 1m-122) and three accessory prototroch cells (1m-211 a sister cell of one of the primary prototroch cells, 1m-1212 and 1m122). The only exception is the 1d quadrant producing only two accessory prototroch cells, due to the migration of the 1d-121 clone out of the episphere (see main text for details). Interestingly, the triplet of the accessory prototroch cells originating from a given quadrant does not associate with the triplet of the primary prototroch cells of the same quadrant, but is rotated by one cell counterclockwise. (C) A detailed scheme of the lineage origin of the apical organ cells.

The apical organ develops from the four cells 1m-111 that form a prominent “apical rosette” in early development, characteristic for the spiral cleavage pattern (Dorresteijn, 1990); Fig. 3A). These cells produce the early differentiating cells of the apical organ (Fig. 3C). A single division of 1c-111 produces the two ampullary cells described previously (Marlow et al., 2014). The two daughters of 1d-111 form the “huge dorsal apical cell” and one of the flask-shaped cells of the apical organ (Tessmar-Raible et al., 2007). The cell 1a-111 buds off the cell 1a-1112 of unknown identity at around 9hpf. The second daughter cell (1a-1111) divides at around 12hpf to give rise to the earliest ChAT-positive cell (1a-11111). Its sister cell (1a-11112) divides multiple times, eventually producing a clone with bilateral symmetry to the clone descendant from 1b-112121 (purple clones in Fig. 4G), providing an example of bilateral clones not related by lineage (see below). The ventral rosette cell 1b-111 shows variable behavior among embryos, from no division (3/6 observed embryos), one division (2/6 embryos) or more divisions (1/6 embryos). The timing of the first division of 1b-111 ranges from ~12 hpf to ~24 hpf. The large nuclear volume and rather low signal resembles the highly proliferative blast cells and suggest a possible proliferation in later development.

**Figure 4:**
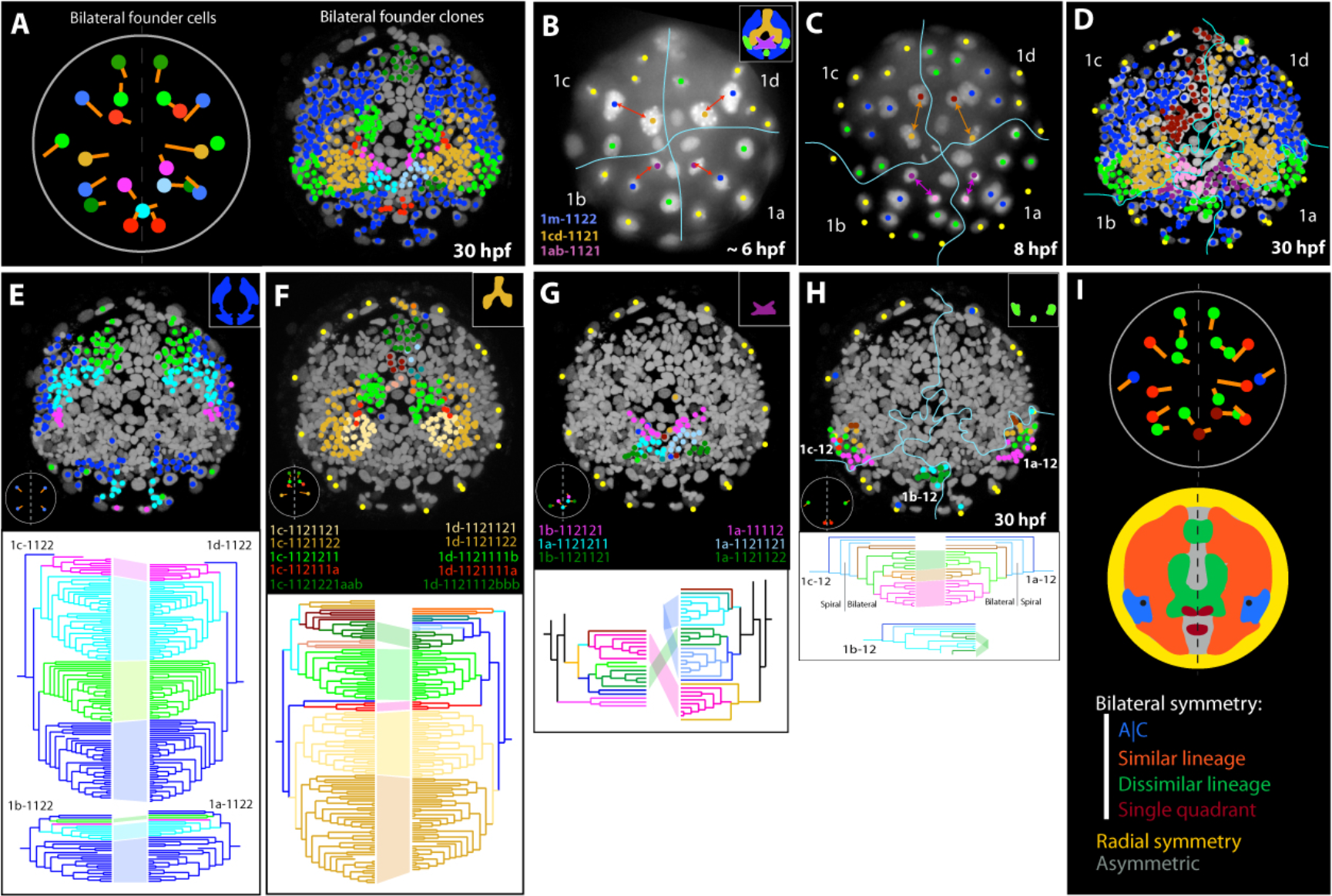
Radically different lineage origin of bilateral founders. (A) The map of bilateral founder cells. The left panel shows the position of bilateral founders the clonal offspring of which is shown in the middle panel. The orange lines represent the division axis between the bilateral founder and its sister cell. Bilateral founder cells originate at different time points during development, summarized in the right panel. (B) The first bilaterally symmetric divisions do not produce bilaterally symmetric clones. (E) The cell divisions of 1m-1122 cells are fully bilateral producing bilateral clones. (F-H) The bilaterally symmetric clonal domains in the medial regions derive from cells with different lineal origin and different lineage depth (for example light green and red clone in F), or magenta clone in G). Note that the bilaterally symmetric light and dark blue clone in G) both originate from 1a quadrant. (H) The cells 1m-12 divide spirally to produce accessory prototroch cells 1m-122 (dark blue) and 1m-1212 (light blue). Subsequent cell divisions within 1c-12 and 1a-12 clone occur in a bilateral mode resulting in fully bilateral domains. (I) The map of bilateral founder cells color-coded for the type of bilateral symmetry. See Supplementary Fig. 4 for more details.

### An array of bilateral founder cells

We next focused on those lineages, for which a transition from the initial rotational to bilateral symmetry was manifested. Previous reports from *Nereis* (Wilson, 1982) and *Platynereis* (Dorresteijn, 1990; Pruitt et al., 2014) identified the first divisions with bilateral symmetry between 7 and 12 hpf, yet, could not track the progeny of these cells at subsequent stages. Using our tracked lineage, we interrogated these cell divisions with regard to their clonal progeny. We defined ‘bilateral founders as cells that would i) have a bilateral counterpart (in position); ii) produce bilaterally symmetrical clonal progeny with similar lineage topology; and iii) appear at roughly the same developmental time point. To our surprise, we identified not only few, but a whole array of pairs of bilateral founders situated on the right and left sides of the *Platynereis* episphere (Fig. 4A). These appear as early as 6 hpf and continue to arise until 18 hpf. The entirety of paired bilateral founders and their bilaterally symmetrical progeny is visible in (Fig. 4A). How do these arise?

The first ‘bilateral’ divisions (i.e., divisions with a bilaterally rather than rotationally symmetrical orientation of spindle poles) take place after less than 6 hpf (Fig. 4B). Specifically, the 1m-112 cells cleave into two bilaterally positioned daughter cells (1m-1121 and 1m-1122). At 8 hpf, these cells continue to divide in a bilaterally symmetrical manner (Fig. 4C). Subsequently, the progeny of these cells divides further and produces all bilateral founders and bilateral founder clones (Fig. 4D; compare with Fig. 4A). Further inspection and comparison of the lineage trees for each pair of left and right founders revealed an unexpectedly diverse and heterogeneous picture for distinct sets of founders – meaning that the transition from rotational to bilateral symmetry involved different strategies for different branches of the lineage tree.

About half of the bilateral founders – the ones located more laterally – are generated in perfect bilateral symmetry, as mirror images on the right and left body sides. This is reflected by a bilaterally symmetrical arrangement of the resulting lateral clones (with dark blue color in Fig. 4D). All descendent lineages show full bilateral symmetry, as is apparent from the equivalent lineage history of right and left counterpart clones (Fig. 4E). Counter-intuitively, the other half of the bilateral founders – the ones located more medially – arise in a very asymmetric manner, which is reflected by the dissimilar outlines of the medial clones on the left and right body sides (with dissimilar colors in Fig. 4D). For each pair of bilateral founders generated by these sublineages (Fig. 4F), the lineage history of the left and right founder is very different. They originate by asymmetric divisions at different branches of the lineage tree and in some cases even differ in the lineage depth. As an example, the offspring of 1c-1121211 and 1d-1121111b (lineage depth difference 1, lineages diverged 4 cell cycles ago) produce a bilaterally symmetric domain (light green clones in Fig. 4F).

Finally, we noted a peculiar difference in how the four initial quadrants 1a, 1b, 1c, 1d contributed to the multiple pairs of bilateral founders. In most cases, founder pairs that were produced from the 1c quadrant on the left side came from the 1d quadrant on the right side, and those produced from the 1b quadrant on the left side came from the 1a quadrant on the right side. In few rare cases, however, pairs of bilateral founders came from the 1a versus 1c quadrants, or resulted from a single quadrant (Fig. 4G-H). The most striking examples are represented by the bilateral clones 1a-1121211 and 1a-1121121 (light and dark blue clone in Fig. 4G) and small clones 1b-12111aa and 1b-121121b (dark green in Fig. 4H) that originate from single quadrants. In summary, the overall contributions of quadrants to bilateral progeny and the processes by which bilateral symmetry is reached are rather complex, forming a “patchwork” of processes schematized in Fig. 4I.

### The conserved apical *six3*, *otx* and *nk21* domains show different lineage behavior

A number of recent studies revealed a conserved role of the homeodomain transcription factors *six3, otx* and *nk21* in the specification of the apical region (Marlow et al., 2014; Steinmetz et al., 2010; Tessmar-Raible et al., 2007). In general, *six3* is expressed most apically, surrounded by a ring of *otx* expression. *Nk21* is expressed in the ventral apical region, overlapping with *six3* and *otx*. Taking advantage of our cellular atlas, we have identified the cells expressing *six3, otx* and *nk21* at 12hpf and compared the clonal progeny of these cells at later developmental time points to the actual expression domains (Fig. 5A-D). This comparison reveals that, while the *six3* and *otx* domains are initially mutually exclusive, *six3* expression later spreads into the *otx* clonal descendants (compare Fig. 5A and Fig. 5D at 24 hpf). In contrast, *otx* expression is turned off in clonal descendants from 20hpf onwards. *Nk21* expression is less dynamic and largely remains expressed in clonal descendants (Fig. 5D).

**Figure 5:**
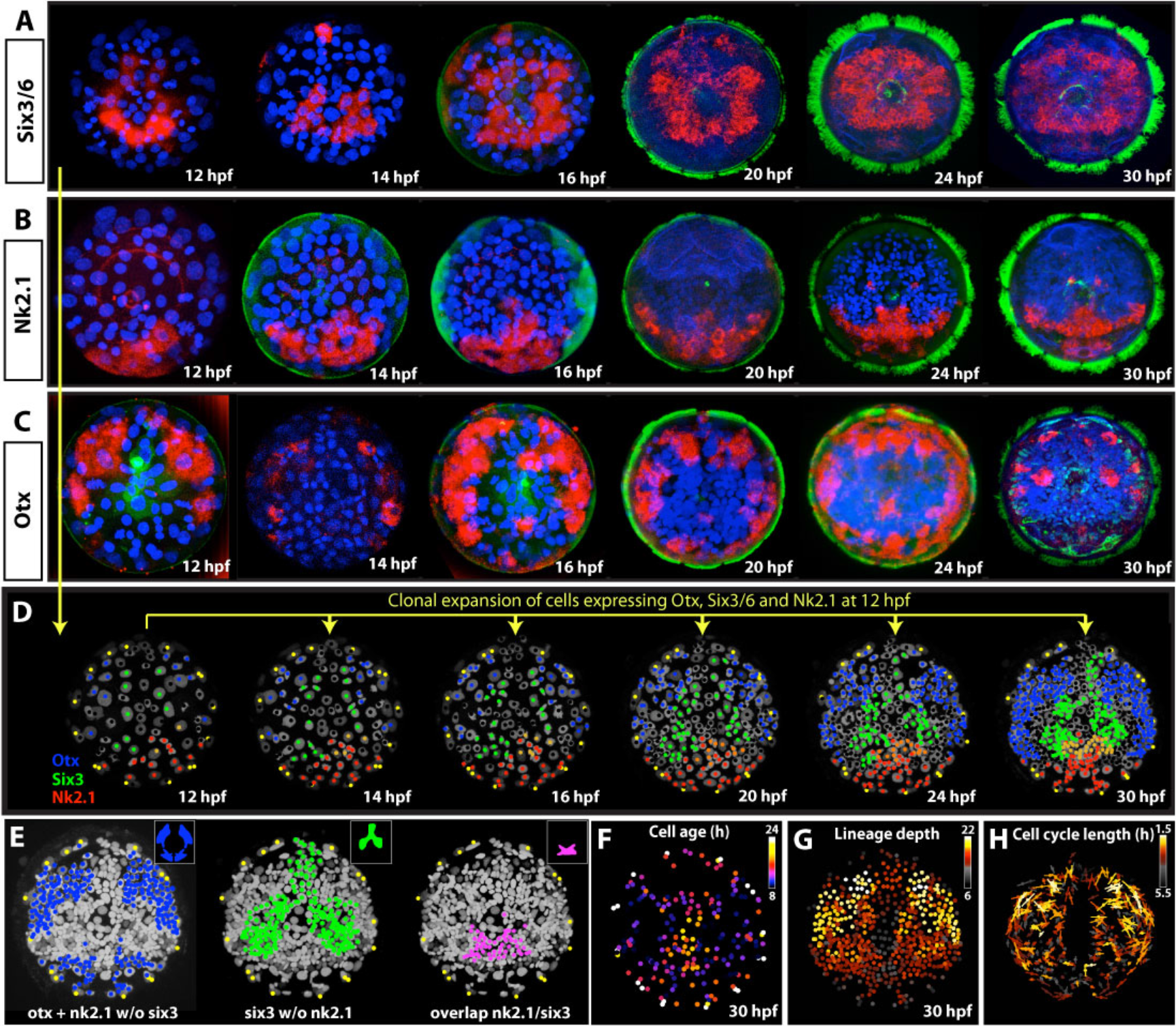
Developmental expression patterns of ancestral early patterning genes. (A-C) Developmental expression of Six3/6 (A), Nk2.1 (B) and Otx (C) between 12 and 30 hpf. (D) The expression of the three genes mapped on the lineage movie at 12 hpf and the clonal offspring of the cells expressing Otx, Six3/6 and Nk2.1 at 12hpf visualized at later stages. (E) Whereas the early Otx and Nk2.1 clonal domain reflects the lateral regions adopting bilateral symmetry very early, the Six3/6 domain encompasses the medial region with later and lineage-unrelated origin of bilateral symmetry. (F) Visualization of cell age (time from the last division) at 30 hpf reveals the prototroch and the apical organ as the earliest differentiating regions of the episphere (compare to expression of the neural markers in Supplementary Fig. 3C). (G) Analysis of the lineage depth (the number of preceeding cell divisions of every cell from 0hpf) identifies the lateral regions as the most proliferative in accordance to the shortest cell cycle length (H).

We set out to characterize the *six3, otx*, and *nk21* further with regard to lineage behavior. At 6 hpf, *otx* is expressed in the 1m-12 primary trochoblast cells (Supplementary fig. 5), which later give rise to the accessory prototroch. At 12hpf, the cells expressing *otx* match the 1m-1122 descendants with few exceptions (Fig. 5C and E), thus including the bilateral founders that produce the set of strictly bilateral clones (compare Fig. 4C). This means, the *otx* domain develops from specific quartets of micromeres, in line with a possible specification by maternal determinants. In addition, the dorsal *otx* region is most proliferative among episphere cells in that it shows the highest lineage depth and the shortest cell cycle length (Fig. 5G-H). Except for the prototroch and accessory prototroch cells, it produces no differentiated cells until 22hpf (whereas the ventral cells 1ab-1122 give rise to the gland cells, Table 1). Cells in this territory differentiate much later, such as the adult eyes (Arendt et al., 2002).

In contrast, at 12hpf *six3* matches the 1m-1121 quartet (Fig. 5A and Fig. 4B), suggesting maternal specification. Yet, the bilateral founders emanating from these cells do not represent quadriradial homologs (and thus are unlikely to be specified maternally). On average, the *six3* cells divide less, and generate several differentiated cells at 22hpf, including the crescent cell (Nr 40 in Table I and Fig. 2F), three ventral ChAT-positive cells (Nr 47, 48 and 50), and one serotoninergic cell (Nr 53). The 12hpf *nk21+* clones are partially co-expressing *otx* and *six3.* They represent the region with highest disorder with regard to bilateral founder cells. Notably, the ancestral patterning genes *Six3, Otx* and *Nk2.1* are absent from the early differentiating apical organ cells that stems from the 1m-111 lineages.

## DISCUSSION

Here we reconstructed the full lineage of the larval episphere in the marine annelid *Platynereis dumerilii*, from spiral cleavage to fully bilateral larval stages, including individual lineages for 62 differentiated cells that make up the larval body. Overall, our data confirm earlier observations that the development of spirally cleaving embryos is highly stereotypic, and extend these observations to early larval stages. Consistent with this, we find that the cell lineage of early differentiating cells is highly invariant.

To relate the *Platynereis* lineage to embryonic and larval stages, we constructed a gene expression atlas for several embryonic and larval stages, for 23 genes with known roles in developmental specification and cellular differentiation. Taking advantage of this multi-stage expression atlas, we have explored how gene expression and early cellular differentiation relate to the transition from rotational to bilateral symmetry.

Lineage tracking, concomitant with the mapping of gene expression and identification of differentiated cells in the consensus lineage tree, is part of ongoing efforts (Achim et al., 2018; Achim et al., 2015; Tomer et al., 2010; Vergara et al., 2017) to resolve and understand *Platynereis* development at single cell level. The comparison of these resources to similar pioneering efforts in other developmental models (e.g. (Du et al., 2015; Santella et al., 2016; Stach and Anselmi, 2015; Tassy et al., 2010), will be especially rewarding for understanding the evolution of development at cell type level.

### Highly complex transition from rotational to bilateral symmetry

Our analysis of the full lineage until 30 hpf has allowed an in-depth investigation of the transition from the embryonic spiral cleavage pattern with rotational symmetry to the bilateral symmetry of the swimming trochophora larval stage. As anticipated by Wilson (Wilson 1892), we find that the bilaterally symmetrical parts of the larval body emerge from so-called bilateral founders. However, the generation of these founders from within the spiral cleavage pattern is surprisingly diverse (Fig. 4I), in that the initial quartets of micromeres produce bilateral founders in very different ways.

First, and most intuitively, bilateral founders emerge from two cells of similar (corresponding) lineage in different quadrants, located on the future left and right body sides. This straightforward strategy applies to about half of the bilateral founders (1m-1122) located more laterally in the larval episphere, However, we detected an unexpected difference in how left-right opposing quadrants contribute to these founders. While, in most cases, the left and right cell of a given founder pair stem from the 1a versus 1b, or 1d versus 1c quadrants, a small portion of the episphere bears an A|C bilateral symmetry established by the descendants of the non-adjacent A and C quadrants (blue regions in Fig. 4I). Remarkably, while the A|C bilateral symmetry is less frequent in *Platynereis* and in other annelids such as *Capitella* (Meyer et al., 2010), it has shown to be predominant in the mollusks *Ilyanassa* and *Crepidula* (Chan and Lambert, 2014; Hejnol et al., 2007).

As a second strategy, we discovered sets of bilateral founders that emerge from two cells of dissimilar (non-corresponding) lineage in left-right opposing quadrants (green regions in Fig. 4I), involving non-bilateral cell divisions at non-related positions within the lineage tree topology (Fig. 4F-H). Even more intriguing, we also observed “single quadrant bilateral symmetry”, where two symmetric clones originate both from the same quadrant (brown regions in Fig. 4I). These findings contradict the initial assumptions (Wilson, 1892) that, as observed for the 2d and 4d somatic descendants in the larval hyposphere, strictly bilaterally symmetric divisions should establish the bilaterally symmetric portions of the larval body.

### Geometrical constraints may explain the different lineage behavior of bilateral clones

Simple geometrical rules may explain the strikingly different lineage behavior of bilateral clones, as illustrated in (Fig. 6). In a four-fold radially (or rotationally) symmetric arrangements (color coded in Fig. 6B), the diagonals represent the only areas naturally exhibiting bilateral symmetry. These could therefore adopt bilateral behavior automatically, without any need for rearrangement or re-specification. Indeed, our analysis of the rotational-to-bilateral symmetry transition revealed that the clones positioned on the diagonals are the first ones to adopt bilateral behavior (Fig. 6A). In other regions of the spiral cleavage pattern, cells of corresponding lineage in left-right opposing quadrants do not automatically come to lie in mirror-image positions. Instead, in the more medial regions between left and right diagonals, left-right mirror image coordinates are of dissimilar origin within opposing quadrants. We propose that this is reflected by highly asymmetric lineage origin of the bilateral founder cells in these regions.

**Figure 6:**
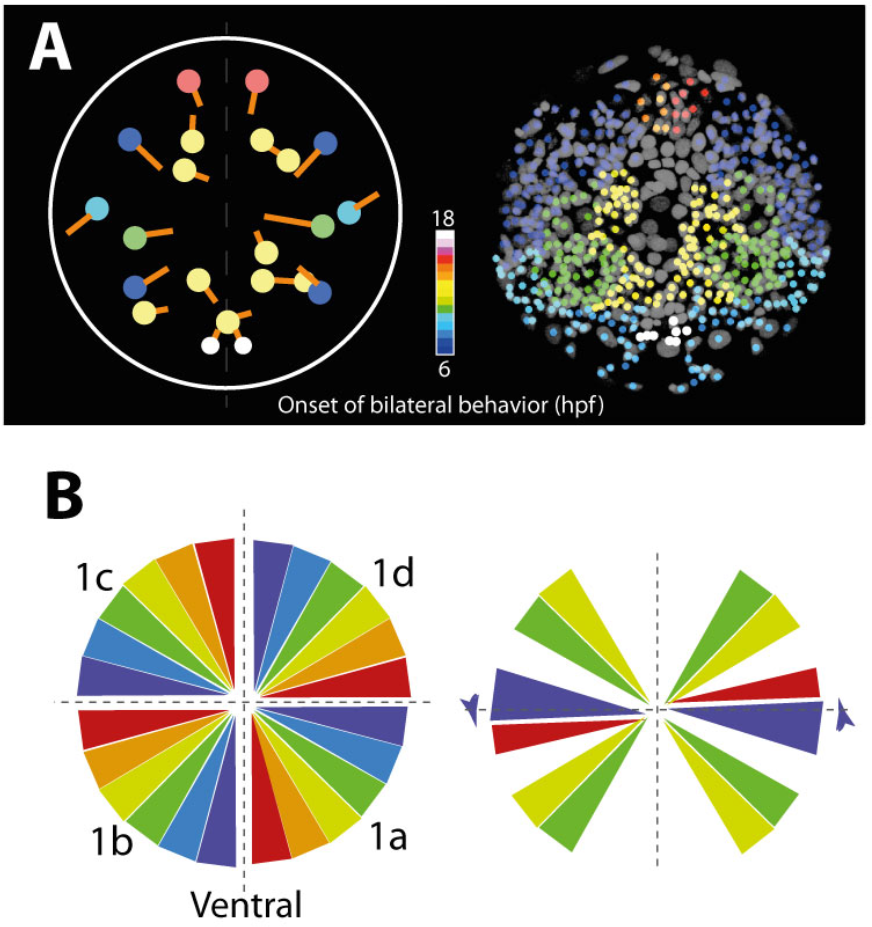
Geometrical constraints in spiralian development. (A) The timing of the transition of blastomeres to bilateral behavior. (B) A theoretical diagram showing four radially symmetrical quadrants (left-hand side) and a subset of areas more susceptible to gain bilateral behavior: The yellow/green areas lying on the diagonals are both radially and bilaterally symmetrical; the purple/violet areas at the interface between 1c/1b or 1d/1a quadrants can become bilateral by expansion of either B and D region (purple triangles) or A and C region (violet triangles). This is indeed a case, where the 1c-12 cell expands ventrally while the 1a-12 expands dorsally to produce bilaterally symmetric domains (blue areas in B). Note that the region along the dorso-ventral midline is conceptually the most difficult to convert into bilateral symmetric structure.

Bilateral symmetry can be also achieved by expanding the areas at the interface between dorsal and ventral quadrants (purple triangles and arrow in Fig. 6B). Dorsal expansion of the ventral quadrant on one side and ventral expansion of the dorsal quadrant on the other side of the embryo produce bilateral symmetry. Indeed, the A|C bilateral symmetry is achieved by such an expansion (light green clones in Fig. 4D). With the above-mentioned assumption of these geometrical constraints it is easy to envision the difficulty in adopting bilateral symmetry within the midline region observed during episphere development. Hence, the axis of bilateral symmetry itself is mostly devoid of cells adopting bilateral behavior and only forms small bilateral clones that emerge at later larval stages.

### A signaling centre in the plane of bilateral symmetry?

Taken together, these observations suggest that there is no obvious relationship between the embryonic cell lineage of the spiral cleavage and the future bilateral symmetry of the larval body. Rather, the common theme of all cells adopting bilateral behavior appears to be their symmetric position with regard to the bilateral axis. This suggests involvement of a signaling source positioned in the plane of the bilateral symmetry axis. An obvious candidate for the signaling center is the cell 2d and its descendants, positioned in the anterior part of the dorsal hyposphere on the axis of the bilateral symmetry. These cells are well known for their organizing potential of *Platynereis* trunk (Pfeifer et al., 2014) and the deletion of the 2d cell in *Capitella* leads to loss of bilateral symmetry in the head (Amiel et al., 2013).

Interestingly, the regulative potential of the D quadrant does not seem to be limited to the C|D-A|B bilateral symmetry, but might also contribute in establishing the A|C bilateral symmetry, as demonstrated by its involvement in specification of the A and C quadrant-derived eyes in *Ilyanassa* (Sweet, 1998).

### Non-bilateral origin of early differentiating cells

The step-wise and complex transition from spiral to bilateral symmetry is also reflected in the different lineage behavior of early differentiating cells, both spatially and temporally. For example, the earliest differentiating prototroch cells have a spiral origin and the equally early-appearing cells of the apical organ are formed by non-symmetrical clones. Earlier work in *Platynereis* (Dorresteijn and Graffy, 1993) and early cell dissociation experiments in *Nereis* (Costello, 1945) pointed to a high degree of cell-autonomous differentiation for these cells via the inheritance of maternal determinants. Several studies in mollusks (Damen et al., 1994; Lambert and Nagy, 2002; Rabinowitz and Lambert, 2010) and in *Platynereis* (Pfeifer et al., 2014) have demonstrated that mRNA segregation into specific blastomeres during the cleavage plays a crucial role in cell autonomous specification.

The spiral and bilateral division patterns co-exist for a certain period, with the first bilateral divisions commencing at ~6 hpf while the last spiral divisions of accessory prototroch cells happen at ~8hpf. Consistently with the notion that the zygotic expression is necessary for the first bilaterally symmetric division in the leech *Helobdella* (Schmerer et al., 2013), we did not observe any bilateral behavior before the onset of zygotic transcription (Chou et al., 2016).

### Six3/6 and Otx specify medial versus lateral bilateral founder cells

Interestingly, the early differentiating cells of the prototroch and the apical organ that are potentially specified via maternal determinants do not express the regional specification transcription factors of the bilateral head regions, Six3/6, Otx and Nk2.1 (Marlow et al., 2014; Tosches and Arendt, 2013). In contrast, *six3, otx* and *nk2.1* expression encompasses all bilateral founders that arise from the 1m-1121 and 1m-1122 micromeres, and thus all lineages of subsequently differentiating cells with bilateral symmetry – at least transitorily. Among these, *six3* expression labels the more medially located 1m-1121 founders that are of different lineage in opposing quadrant, whereas *otx* labels the more lateral bilateral founders that stem from 1m-1122 micromeres, with similar lineages between quadrants. The *six3+* founders give rise to a bilateral subset of cholinergic neurons that differentiate early in the larval brain. In contrast, the *otx+* lateral founders proliferate heavily during later stages and will differentiate much later, into adult eyes and optic lobes (Arendt et al., 2002).

From these observations some important insights emerge with regard to the possible autonomous specification of larval lineages via maternal determinants. First, given that the *six3+* and *otx+* cells show corresponding lineages in each quadrant (1m-1121 and 1m-1122 micromeres), it is possible that the expression of these transcription factors is regulated autonomously (pending experimental validation). At the same time, however, the dissimilar and asymmetric sublineages of all cells differentiating from the *six+* founders precludes a role for maternal determinants and instead suggests conditional specification of the cell fates that emerge from the *six3+* micromeres.

Future integration of our lineage data with single-cell expression data mapped onto the expression atlases constructed for reference embryonic and larval stages, will allow the identification of candidate signals and receptors, as well as the gene regulatory networks establishing bilateral symmetrical behavior and cell fates in each of the founder lineages.

## MATERIALS AND METHODS

### Animals

The larvae of *Platynereis dumerilii* were obtained from the breeding culture at EMBL Heidelberg.

### Injections and time-lapse imaging

The injections of H2A-mCherry and mYFP were performed as described previously (Lauri et al., 2014). For tracking axonal projections, LifeAct-EGFP mRNA (Benton et al., 2013) was injected into a given blastomere of embryos injected previously at one-cell stage with H2A-mCherry mRNA.

The injected embryos were kept in filtered seawater at 18°C until the desired developmental stage was reached. Selected embryos were then transferred in ~2μl of sea water into 40°C warm 0.8% low-melting agarose (A9414, Sigma-Aldrich), briefly mixed by pipetting up and down and quickly transferred in ~20μl agarose to the microscopy slide with 150 μm spacer on each side (3 layers of adhesive tape Magic™Tape, Scotch^®^). Before the agarose fully solidified (within ~15 seconds) the embryos were covered by a coverslip and oriented to the apical position for imaging. Seawater was added from the side of the slide to entirely fill the slide chamber. To avoid drying out, the coverslip was sealed using mineral oil. The embryos were imaged using a Zeiss Axio Imager.M1 fluorescent microscope or Leica TCS SPE confocal microscope with 40× oil immersion objective with time resolution 20 min (Movie 3), 15 min (Movie 8, 10 and 11), 6 min (Movie 1) and 12 min (Movie 2).

### Tracking and comparing the cell lineage across multiple embryos

The live-imaging movies were manually tracked using a custom-made tracking macro in ImageJ/FiJI (Schindelin et al., 2012). We used the nuclei count of the episphere in embryos precisely fixed at several time points to calibrate the developmental time in the movies. Due to a high density of nuclei at later stages, we were able to reliably track until about 32hpf. We use the standard spiralian nomenclature of the cells according to (Conklin, 1897).

After 6hpf, even for non-spiral cell divisions we use the index 1 for the more anterior and index 2 for the more posterior daughter cell until around 10hpf. After 10hpf, we use indices “a” and “b” instead of “1” and “2” and without any added information. To compare the cell lineage across different embryos, we developed a simple algorithm (Supplementary Fig. 1F) automatically identifying corresponding cells in each tracking dataset and highlighting the differences (see also Supplementary materials and methods).

### Whole-mount mRNA in situ hybridization

The mRNA *in situ* hybridization was performed as described in (Tessmar-Raible et al., 2005) with following modifications: For developmental stages earlier than 12hpf, the embryos were washed twice 4 minutes with TCMFSW (Schneider and Bowerman, 2007) prior to fixation. For developmental stages younger than 24hpf, the embryos were acetylated: After the digestion in proteinase K and two washes with freshly prepared 2mg/ml glycine in PTW (1× phospate-buffered saline with 0.1% Tween-20), the embryos were incubated 5 minutes in 1% triethanolamine in PTW, then 3 minutes in 1% triethanolamine with 0.2% acetanhydride followed with 3 minutes of 0.4% acetanhydride in 1% triethanolamine. The prehybridization, hybridization and SSC washes were performed at 63°C. The hybridization mixture: 50% Formamide (Sigma-Aldrich, F9037), 5× SSC pH4.5, 50 μg/ml Heparin (Sigma-Aldrich, H3149), 0.025% Tween-20 (Sigma-Aldrich, P9416), 50 μg/ml Salmon Sperm DNA (Sigma-Aldrich, D9156), 1% SDS. The DIG-labeled antisense mRNA probes: ChAT, Elav (Denes et al., 2007); Syt, TPH, Phc2, Nk2.1, (Tessmar-Raible et al., 2007); VACht (Jekely et al., 2008); Otx (Arendt et al., 2001); Six3/6 (Steinmetz et al., 2010); VGlut (Tomer et al., 2010).

